# Cooperativity and Communication between the Active Sites of the Dimeric SARS-CoV-2 Main Protease

**DOI:** 10.1101/2025.07.31.667456

**Authors:** Sarah N. Zvornicanin, Ala M. Shaqra, Julia Flynn, Lauren E. Intravaia, Heidi Carias Martinez, Weiping Jia, Devendra Kumar Gupta, Stephanie Moquin, Dustin Dovala, Daniel N. Bolon, Brian A. Kelch, Celia A. Schiffer, Nese Kurt Yilmaz

## Abstract

The coronaviral main protease (M^pro^) is essential for the replication of the virus, and has been the subject of various biochemical, structural and enzymatic studies, as well as a drug target against SARS-CoV-2 infections. SARS-CoV-2 M^pro^ is known to be active as a dimer, with the N terminus of one protomer completing a key active site pocket of the other protomer. Despite apparent cooperativity in catalytic activity, how the two distal active sites in the dimer communicate and might be modulating binding and/or catalysis at the other remain to be clarified. Here, we have investigated the interplay between cooperativity, dimerization, and substrate cleavage in SARS-CoV-2 M^pro^ through a combination of enzymatic assays, crystal structures, and protein characterization. To disentangle the contribution of each active site to the observed enzymatic activity, we developed a cleavage assay involving heterodimers of active and inactive (C145A or inhibitor-bound) monomers. Strikingly, we found that heterodimerization increased cleavage efficiency per active monomer. Additionally, we mapped a network of critical residues bridging the two active sites and probed this network through engineered mutations. By dissecting the cooperativity and communication between the active sites, we provide new insights into the M^pro^ reaction cycle and functional significance of its dimeric architecture.

## Introduction

The replication of coronaviruses, including SARS-CoV-2, relies on the enzymatic function of the viral main protease (M^pro^), which cleaves viral polyproteins to release structural and non-structural proteins required for viral assembly and maturation. As a result, M^pro^ is a key target for COVID-19 antiviral therapies. SARS-CoV-2 M^pro^ is known to function as a dimer with two distant active sites, while the monomeric form is inactive and lacks a properly formed active site.^1–11^ Mutations that disrupt dimerization impair enzymatic activity, highlighting the importance of stabilizing the functional dimer.^7,10,12^ M^pro^ exhibits apparent positive cooperativity in peptide cleavage assays, as indicated by a Hill coefficient greater than one.^13–16^ However, positive cooperativity in ligand binding (thermodynamic allostery) does not explain M^pro^’s observed behavior, which led to proposals of negative cooperativity in non-covalent ligand binding^13,17^ as well as half-site reactivity (that only one monomer of the dimer is active) in the very closely related SARS-CoV-1 M^pro^.^18^

While evidence suggests cooperativity and allosteric communication between the two active sites, the mechanistic basis for this allostery is yet to be established. The dimeric M^pro^ is symmetric in the absence of bound ligands, while binding at one of the active sites would break the symmetry and could allosterically induce rearrangements at the other active site. M^pro^ recognizes and cleaves substrates of diverse amino acid sequences.^19,20^ Our cocrystal structures of SARS-CoV-2 M^pro^ (inactive C145A variant to prevent cleavage) with peptides corresponding to 9 viral cleavage sites revealed both active sites to be symmetrical and fully occupied with the same ligand binding mode.^19^ While M^pro^ rearranges loops and regions around the active site to accommodate various ligands, this rearrangement is virtually the same between the two protomers (in crystal structures with more than one chain in the asymmetric unit). Likewise, the cocrystal structures of active M^pro^ with covalent or non-covalent inhibitors display fully occupied virtually identical active sites in both protomers.^2,21–23^ However, the enzyme’s reaction cycle likely involves an intermediate state with a single-occupied active site and this initial binding might modulate binding and/or cleavage at the other active site. How one protomer might affect the other and the mechanism of communication between the two active sites remain to be characterized.

The gaps in knowledge about the kinetic steps in the reaction cycle of M^pro^ lead to several questions: How is binding at one active site communicated to the other to yield positive cooperativity? Does each monomer contribute equally to cleavage, and do both active sites need to be occupied for cleavage? How does M^pro^’s cooperativity increase cleavage rate, and ultimately, why is M^pro^ a dimer?

To probe these questions, we have taken a biochemical and structural approach to characterize cooperativity, dimerization, and substrate cleavage. To disentangle the contribution of each active site to the observed enzymatic activity, we developed an innovative cleavage assay involving heterodimers of active and inactive (C145A or inhibitor-bound) M^pro^ monomers. Strikingly, we found that heterodimerization increased cleavage efficiency per active monomer. Additionally, we mapped a network of critical residues bridging the two active sites, hypothesizing that their structural linkage facilitates inter-monomer communication. Variants with mutations at these network residues were assessed for dimerization, thermostability, enzymatic activity, and structural changes through 8 new crystal structures. By dissecting the intricate relationship between M^pro^’s active sites, we provide new insights into the functional significance of its dimeric architecture.

## Results

### Heterodimerization with inactive M^pro^ increased substrate cleavage per active monomer

SARS-CoV-2 M^pro^ functions as a dimer and has been reported to exhibit positive cooperativity,^13–16^ which we confirmed in a cleavage assay using a fluorophore-labeled peptide (see Methods). For WT homodimers, we measured a *K*_m_ of 11.4 ± 0.6 mM and catalytic efficiency of 2.9 ± 0.2 mM^−1^s^−1^ (**Figure S1**).

To probe the observed cooperativity and inter-monomer communication, we designed an assay where only one of the monomers is catalytically active. In this assay, we ensured that virtually all the observed fluorescence signal comes from heterodimers: a dimer composed of one monomer of wild-type (WT) M^pro^ and one monomer of a catalytically inactivated (C145A) M^pro^ (**Figure 1a**). To achieve this, an excess amount (5 mM) of inactive M^pro^ was mixed with a relatively small amount (20 or 40 nM) of active M^pro^ such that after monomer exchange >99% of active monomers were present in WT/C145A heterodimers. During equilibration, the substrate cleavage activity was measured periodically which increased steadily for about 2 hours of incubation before reaching ~1.5-fold of the starting value (**Figure 1b**). Thus, the WT/C145A heterodimers were more active compared to WT dimers, at the same concentration or number of catalytically active monomers (**Figure 1**). In the WT/C145A heterodimers, because the inactive monomer cannot cleave but can bind substrate, we concluded that binding at this active site enhanced the catalytic activity at the other active site. Activity increase upon WT/C145A heterodimerization suggests that active sites contribute non-additively to the total cleavage rate in the WT homodimers.

**Figure 1.**
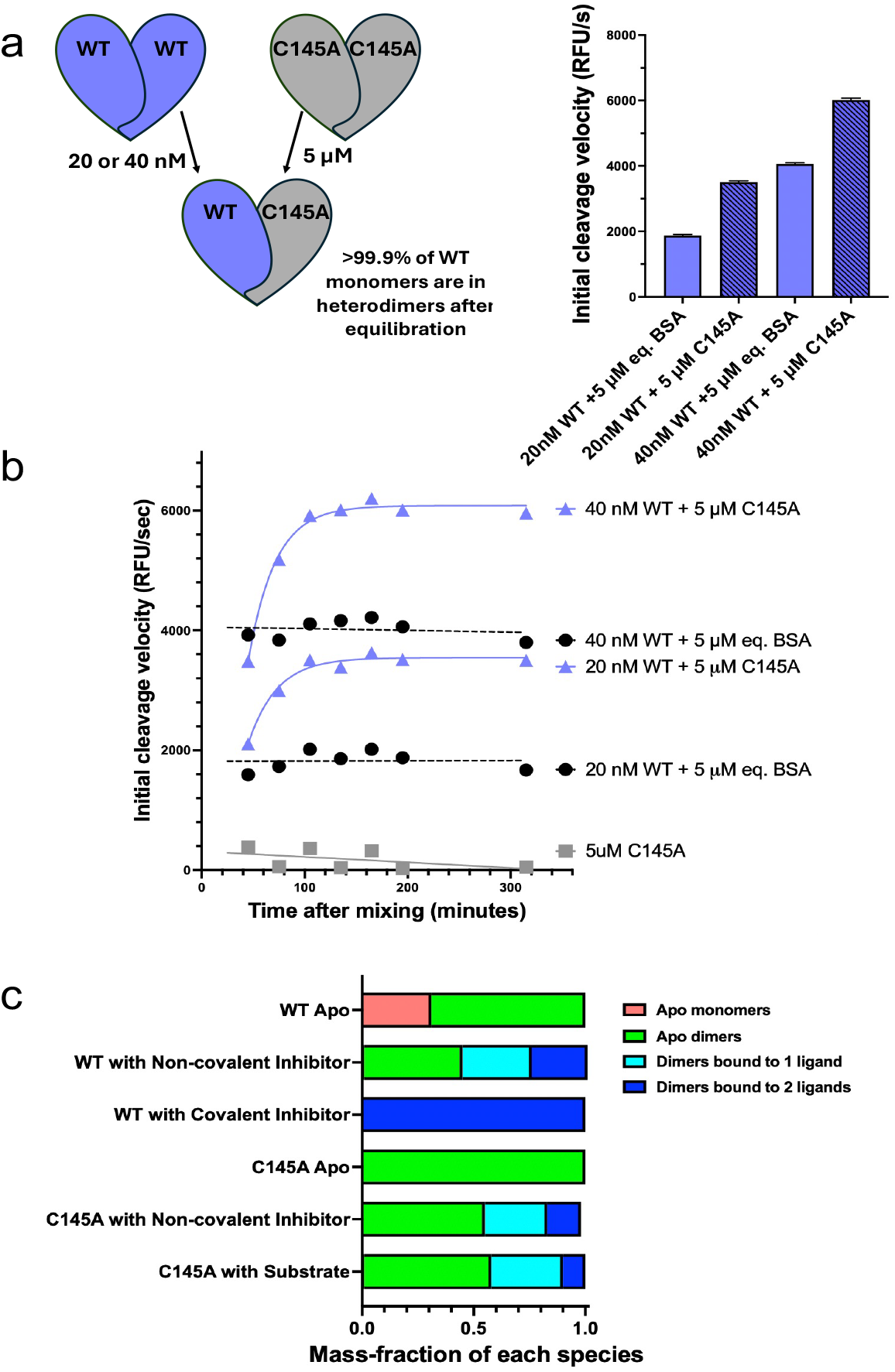
SARS-CoV-2 M^pro^ heterodimerization with inactive monomer, and ligand-induced dimerization. The rate of substrate cleavage during heterodimerization with catalytically inactive C145A M^pro^ (**a**) final observed cleavage rate after equilibration, and (**b**) change of cleavage rate over time after addition of excess amount of inactive M^pro^. BSA was added to the control to ensure equal protein concentration. Error bars indicate standard error of the mean from three replicates. (**c**) The fraction of monomeric and dimeric states assessed by native mass spectrometry. The non-covalent inhibitor (Moonshot X11612), reversibly covalent inhibitor (GC376) and substrate (peptide sequence representing nsp4-nsp5) were used as ligands.

To further probe the mechanism of cleavage rate increase, we formed heterodimers with a constitutively ligand-bound M^pro^ monomer. First, WT M^pro^ was inhibited with saturating concentrations of irreversible covalent Michael acceptor inhibitor N3^22,24^ after which the unbound N3 was removed to yield completely inhibited dimers (see Methods). Purified N3-bound M^pro^ was mixed with a relatively small amount of active WT M^pro^, as in the C145A experiments, to yield WT/WT:N3 heterodimers.

With the WT/WT:N3 heterodimers, equilibration was much slower but the substrate cleavage activity increased over time as with the WT/C145A heterodimers (**Figure S2**). After 72 hours of incubation, M^pro^ cleavage activity per active monomer almost doubled indicating that constitutive binding strongly enhanced the apparent enzymatic activity. The longer equilibration time indicates that inhibitor binding substantially decreased the rate of monomer exchange. The increase in substrate cleavage rate when WT monomers exchanged into heterodimers suggests that binding (but not cleavage) at one active site increases the catalytic activity of the other monomer, leading to the observed positive cooperativity of M^pro^.

### Dimerization and ligand-binding to WT and C145A SARS-CoV-2 M^pro^

We previously measured SARS-CoV-2 M^pro^ to have a dimerization *K*_d_ of 0.35 μM^25^, which indicates sampling of monomeric and dimeric species under our assay conditions. We used native mass spectrometry to quantify the dimer fraction and characterize ligand-induced dimerization of M^pro^ in response to substrate, non-covalent inhibitor (Moonshot X11612), and covalent inhibitor (GC376) binding (see Methods). At 5 mM, about 30% of WT M^pro^ was monomeric in the apo state, which completely dimerized upon addition of either covalent or non-covalent inhibitor (**Figure 1c and Figure S3**). While M^pro^ displayed complete ligand-induced dimerization, unexpectedly, the population consisted of a mixture of apo, singly bound and doubly bound dimers. With the non-covalent inhibitor, 44 ± 1% was apo dimers, 31 ± 1% was dimers bound to a single ligand, and 25 ± 1% was dimers bound to two ligands. When the covalent inhibitor GC376 was added to WT M^pro^, the population shifted entirely to doubly bound dimers.

Lastly, we confirmed the ability of catalytically inactive C145A variant to bind substrate and noncovalent inhibitors. In the absence of ligand, the C145A variant was fully dimeric, indicating a lower *K*_d_ of dimerization compared to WT M^pro^. The catalytically inactive C145A variant also yielded a mixture of apo, singly bound, and doubly bound dimers with slightly lower (~10% less) fraction of doubly bound dimers with either substrate or non-covalent inhibitor (**Figure 1c**). Thus, albeit lower affinity, the C145A variant is able to dimerize and bind substrate, in agreement with the hypothesis that substate binding at the inactive monomer enhances the substrate cleavage of the active monomer in the WT/C145A heterodimerization experiments.

### Identification of a network connecting the two active sites of SARS-CoV-2 M^pro^

Given our evidence that binding at one active site influences the activity of the other, we sought to identify the inter-monomer connectivity and the molecular network facilitating communication between the two active sites. We previously reported a comprehensive fitness landscape of SARS-CoV-2 M^pro^ where all possible point mutations were introduced to the enzyme to measure the impact on function^26^. This analysis identified both positions tolerant to substitutions as well as those that cannot be mutated without disrupting enzymatic activity. A particular region of interest was a network of residues bridging the two active sites, which exhibited high sensitivity to mutation. The same residues were positionally conserved among our cocrystal structures with substrate peptides bound to C145A M^pro 19^ (**Figure 2a**) (see Methods). The top 20 most consistently positioned residues were compared with the 24 positions within M^pro^ that demonstrate low mutational tolerance, defined as at least 17 out of the 19 possible amino acid substitutions yielding null-like function^26^. Mapped onto the structure of SARS-CoV-2 M^pro^, the two sets of residues highlight a bridge between the two active sites, and show six shared residues: S10, S113, G146, S147, G149, and H163. We selected three positions from these (S10, S113, and S147) and added L115 due to its incorporation in a beta-strand (residues 113-117) supporting the interface residues (**Figure 2b**). The evolutionary conservation at these residue positions across all coronavirus genera also supports their functional importance (**Figure 2b inset**). Positions 10, 115 and 147 are highly conserved, and position 113 is serine, threonine, or asparagine, which are residues with similar polarity and hydrogen bonding potential. Overall, these residues, which are located between the two active sites of M^pro^, are functionally essential.

**Figure 2.**
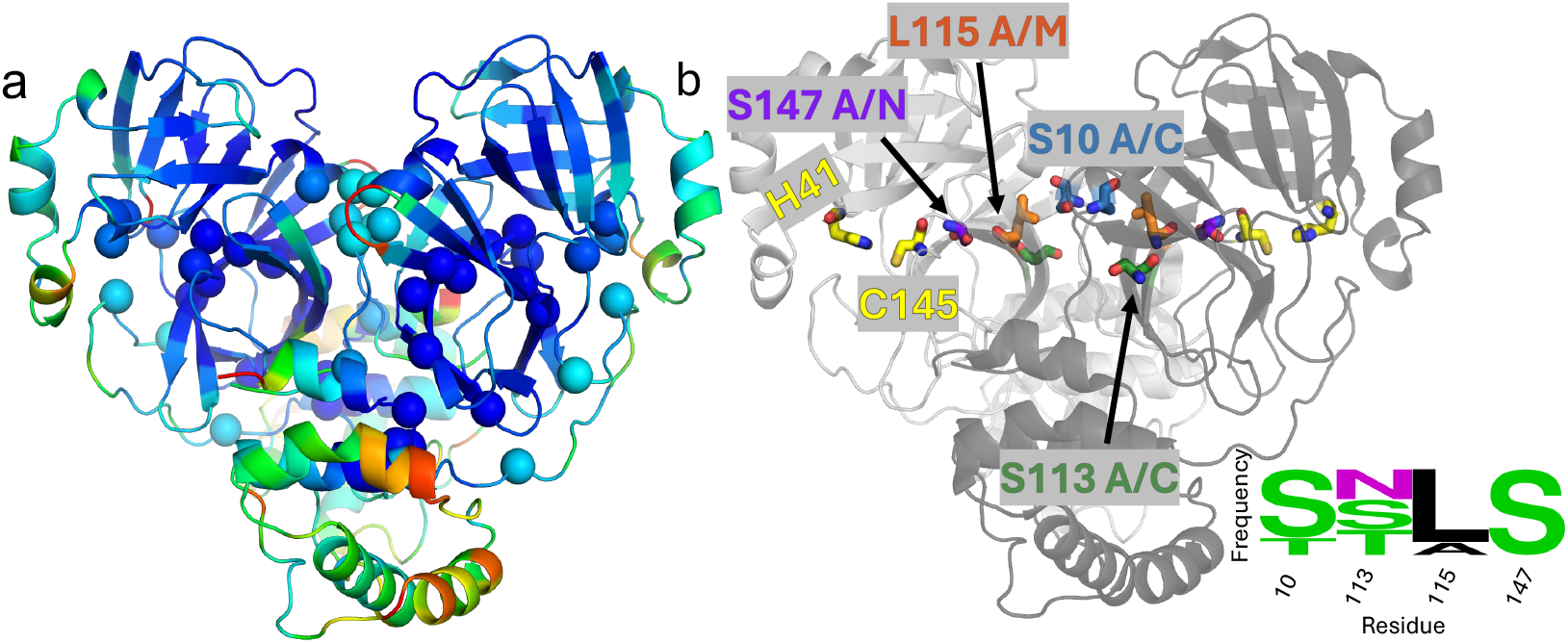
Identification of network residues bridging the active sites in SARS-CoV-2 M^pro^ structure. (a) Residues colored by positional conservation in cocrystal structures bound to one of nine substrates, blue to red for increasing variability. Spheres indicate positions that are intolerant to substitutions and thus essential for function^26^. (**b**) Position of network residues selected to probe via mutations, S10A/C (blue), S113A/C (green), L115A/M (orange), and S147A/N (purple), are highlighted as sticks on the dimer of SARS-CoV-2 M^pro^. Catalytic residues are colored yellow to indicate the active sites, and monomers can be distinguished by shade of grey in the cartoon representation. Sequence logo represents residue conservation in a list of 20 M^pro^ sequences across alpha, beta, delta, and gamma-coronaviruses.

### Cocrystal structures of network variants reveal minor local perturbations

To assess the functional importance of the network residues bridging the two active sites of SARS-CoV-2 M^pro^, we engineered eight network mutations. For each of the selected four positions, we introduced an alanine, and a mutation that was intended to partially rescue WT-like interactions (**Figure 3**).

**Figure 3.**
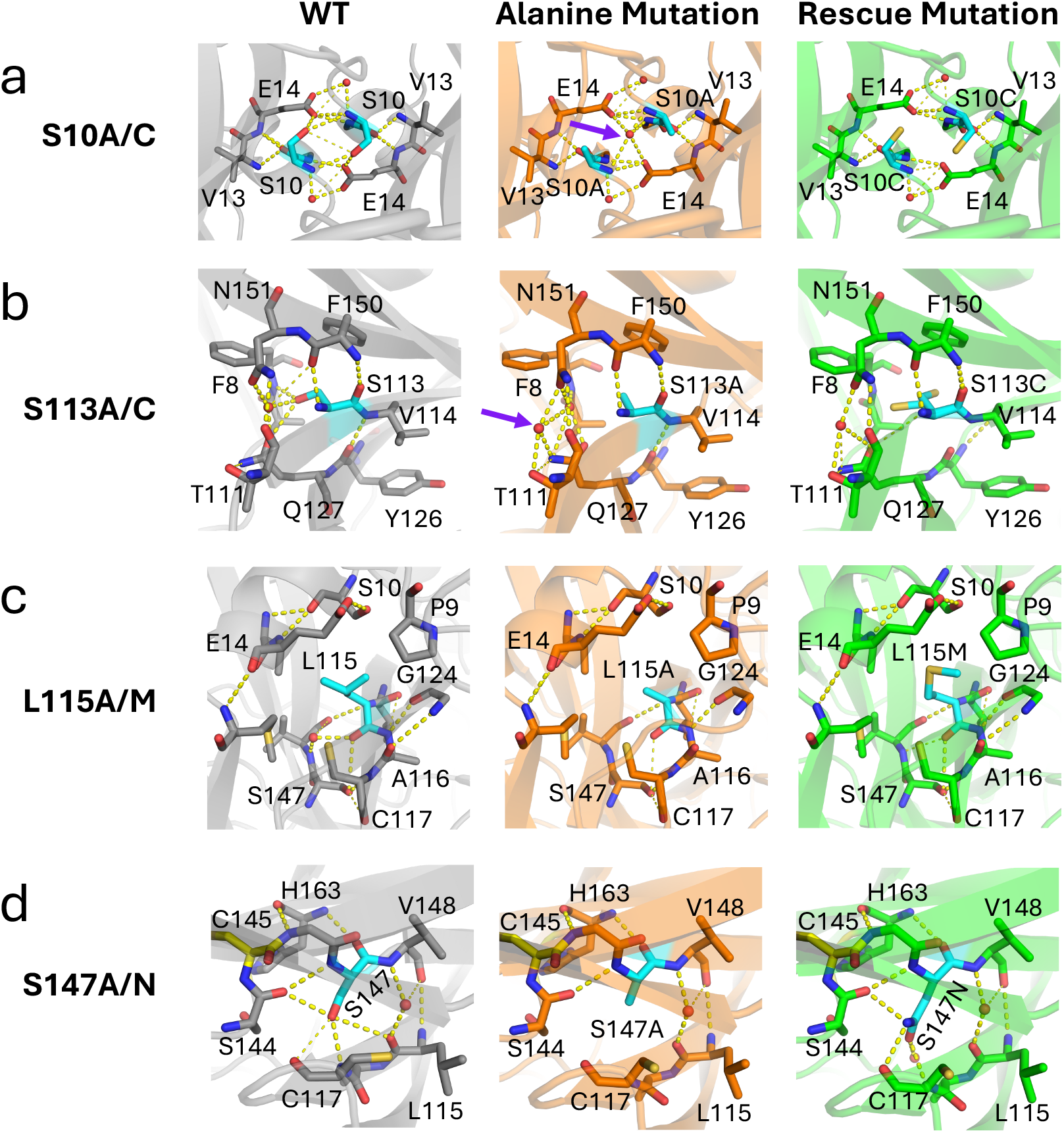
Crystal structures of SARS-CoV-2 M^pro^ network variants. The local structure around the site of mutation is shown for WT (PDB ID: 8DSU) and the mutants for (a) S10 (b) S133 (c) L115 and (d) S147. The mutation site is colored cyan. Waters are represented as red spheres, and dashed yellow lines indicate hydrogen bonds. Residues involved in interactions surrounding the mutation site are displayed as sticks.

We determined the cocrystal structures of SARS-CoV-2 M^pro^ S10A, S10C, S113A, S113C, L115A, L115M, S147A, and S147N variants in complex with the inhibitor PF-00835231 to sub-2.3 Å resolution (**Table S1**). The protein complexes formed well-diffracting, small crystals with the inhibitor. Each variant co-crystallized with two monomers composing a single dimer in the asymmetric unit in the P21 space group. Our R-work values for these structures range from 17% to 21% and R-free values range from 21% to 26% (**Table S1**). Both active sites of each variant were fully occupied by the inhibitor, which was bound in the expected conformation observed in WT M^pro^ crystal structure^27^ and with strong electron density.

At the site of mutation, crystal structures of variants demonstrated loss of interactions which were expected during the selection of mutations. Otherwise, there were only subtle alterations in the local structure around the site of mutation. The S10 variants impacted the symmetrical side-chain interactions at the dimer interface, with alanine causing a complete disruption and cysteine partially recapitulating monomer-monomer interactions through alternative polar interactions (**Figure 3a**). S113 exists on a beta-strand close to the dimer interface. The alanine variant lost side-chain interactions with the backbone nitrogen of F8, a coordinated water, and the amide oxygen of Q127 seen in the WT structure. While the S113A mutation disrupted key contacts and introduced a structural water, the dual conformation of cysteine indicates increased flexibility in the S113C variant (**Figure 3b**). The L115 variants had decreased van der Waals interactions with nearby residues in the 120s beta-sheet and the 10s alpha helix, but neither alanine nor methionine substitution altered the backbone conformation (**Figure 3c**). Lastly, substitutions at S147 altered local hydrogen bonding patterns near the catalytic cysteine, with alanine disrupting interactions entirely and asparagine partially restoring them while maintaining overall backbone conformation. S147 is only two residues away from the catalytic cysteine (**Figure 3d**). In the WT enzyme, S147’s hydroxyl group hydrogen bonds with the backbone oxygen of S144. The alanine variant lost this interaction, while the asparagine mutation was indeed able to partially recapitulate WT-like alternative interactions. In both cases, there were no significant differences in C-alpha positions or the backbone relative to the WT structure.

### Substitutions at network residues severely decreased enzymatic activity

Despite only minor perturbations around the site of substitution in the crystal structures, the variant enzymes had very poor enzymatic activity relative to WT (**Figure S1**). The loss in catalytic efficiency (*k*_cat_/*K*_m_) was mostly due to a decrease in the rate constant for the cleavage reaction (*k*_cat_), which was 3–166-fold lower relative to the WT enzyme (**Figure S1**). S10A exhibited the largest loss of catalytic efficiency, with a more than 4-fold decrease in substrate affinity and drastically reduced turnover rate, resulting in almost no detectable catalytic activity under our assay conditions. Both L115A and S147A had catalytic efficiencies less than 3% compared to WT. Overall, the decrease in *k*_cat_ was more pronounced for the alanine variants with the exception of S113A, compared to the corresponding variant with the rescue mutation.

In general, the partial rescue variants had higher substrate affinity and better turnover rates than their alanine counterparts (**Figure S1c**). S10C showed 4-fold improved substrate affinity and 50-fold faster turnover rate than S10A. The most active network variant L115M had unchanged affinity for the substrate (*K*_m_) and 35% catalytic efficiency relative to WT, which was ~10-fold better than L115A. Defying this pattern, S113C exhibited both a higher *K*_m_ and a lower *k*_cat_ than its alanine counterpart S113A, resulting in only 3% catalytic efficiency relative to WT.

The data indicate that mutations within this network severely impair M^pro^’s catalytic efficiency. For the most part, alanine variants had reduced affinity for the substrate and lower turnover rate than the variants designed to structurally recapitulate some WT interactions. The poor enzymatic activity of network variant enzymes highlights the critical role of these residues in maintaining optimal substrate binding and turnover.

In addition to catalytic efficiency, we also explored cooperative behavior by calculating the Hill coefficients for the network variants which were altered compared to the WT enzyme (**Figure S1c**). Hill coefficient values should be interpreted cautiously, since they are challenging to determine with precision and can be even more difficult to interpret^28,29^. Both L115 variants displayed slightly higher Hill coefficients than the WT (1.36 ± 0.08 and 1.41 ± 0.07) while certain variants, including S113A, S147A, and S113C, exhibited Hill coefficients at or below 1.0, potentially indicating some loss in cooperative behavior relative to WT.

### Stability and Dimerization of Network Variants

To investigate whether mutations within the network bridging the active sites disrupt the structural stability of SARS-CoV-2 M^pro^, we compared the melting temperature (T_M_) of each variant to the WT enzyme. WT M^pro^ displayed a T_M_ of 52.9 ± 0.3°C. Interestingly, the catalytically inactivated mutant C145A displayed enhanced thermostability with a T_M_ of 56.8 ± 0.1°C. Most variants exhibited slightly decreased thermostability compared to WT, although the reductions were generally modest (0.8 to 5.4 °C) (**Figure S3**). Except for S147, the rescue mutations were more successful compared to alanine mutations for retaining thermostability. While network variants exhibited reduced thermostability, the measured T_M_ values remain within a range consistent with normal enzymatic function and structural integrity^30,31^. This suggests that global protein destabilization cannot be the reason for the drastic loss in the enzymatic activity of the network variants.

We also assessed dimerization of the network variants, which is essential for SARS-CoV-2 M^pro^ activity.^18^ In the apo state, the network variants exhibited significantly reduced dimerization compared to WT enzyme (69 ± 3% dimeric at 5 μM), with percentages ranging from 7 ± 1% (S147A) to 15 ± 1% (L115M). (**Figure 4a**; see Methods). Even substitutions designed to partially recapitulate wild-type like interactions, such as S10C (12 ± 1%), S113C (10 ± 3%), L115M (15 ± 1%), and S147N (8 ± 1%), showed markedly reduced dimerization relative to WT. Importantly there was no correlation between a variant’s thermostability and either catalytic efficiency (Spearman’s r = −0.08, P> 0.78) or dimerization (Spearman’s r = −0.03, P> 0.93) in the apo state. The correlation between dimerization in the apo state and catalytic efficiency was weak (Spearman’s r= 0.66; P<0.012).

**Figure 4.**
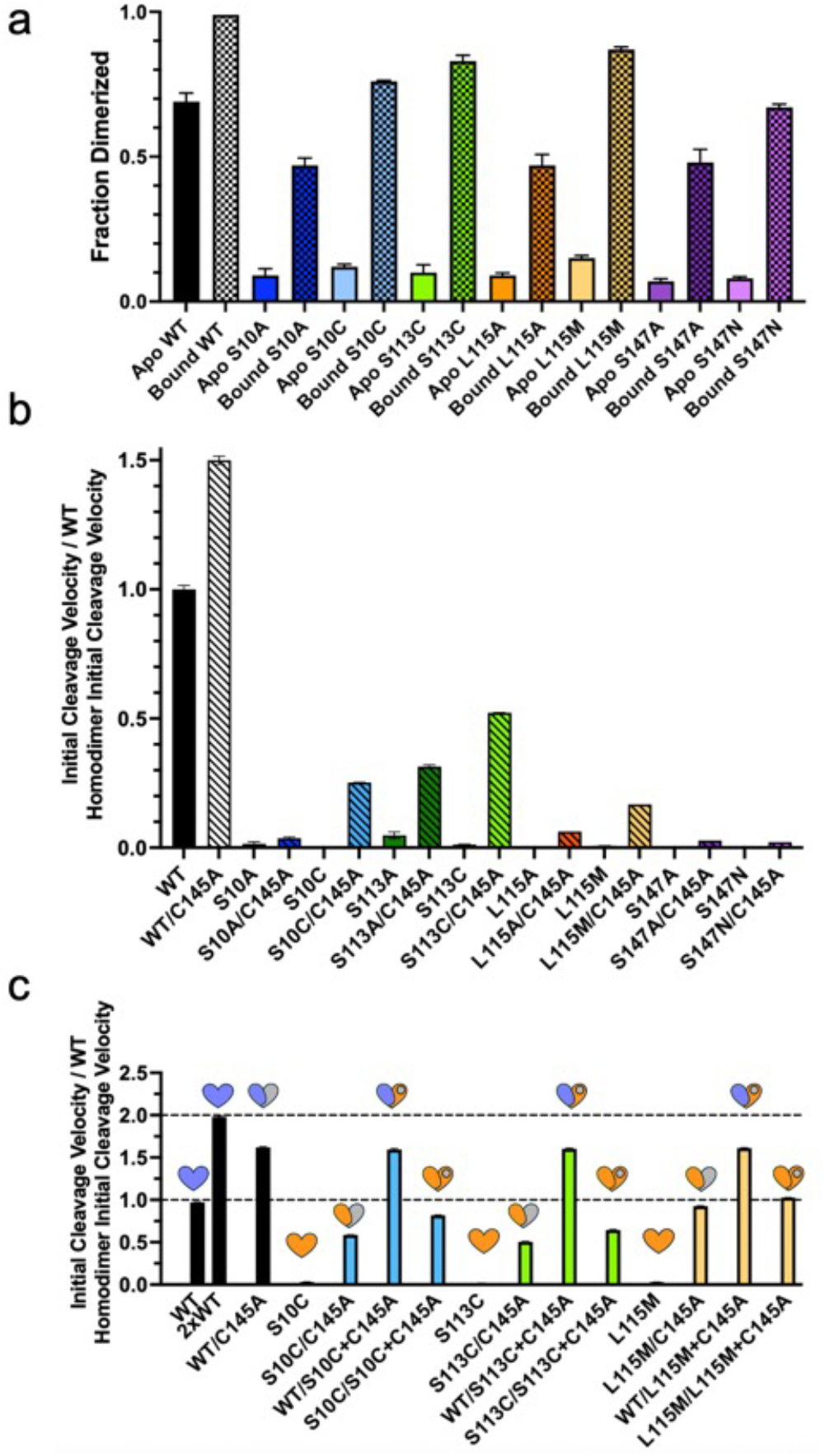
Dimerization and enzymatic activity of SARS-CoV-2 M^pro^ network variants. (**a**) Fraction of dimeric species for apo (solid colors) and inhibitor-bound (hashed bars) M^pro^ variants. (**b**) Rate of substrate cleavage for network variants and their heterodimers with the inactive C145A variant. (**c**) Rate of substrate cleavage for heterodimeric M^pro^ species. The symbols above the bars represent the dimeric species where blue, grey, and orange indicate WT, C145A, and network variant monomers respectively, and gray circle within indicates the presence of the catalytically inactivating C145A mutation. The horizontal dashed lines show activity for WT homodimers at two concentrations for comparison. For cleavage activity assessments of M^pro^ homodimers, BSA was added at a concentration equivalent in mass to that of 5 µM WT M^pro^. Error bars indicate standard error of the mean between three replicates.

Despite the severe loss in dimerization in the apo state, all M^pro^ variants dimerized effectively when bound to a ligand. The non-covalent Moonshot compound X11612 induced dimerization 38% to 72% higher than apo state (**Figure 4a**). Ligand-induced dimerization is known for coronaviral main proteases, including MERS-CoV M^pro^ which is monomeric in the apo state but fully active and dimeric when ligand bound^32,33^.

To further assess ligand-induced dimerization, we expressed and purified three catalytically inactivated network variants: C145A+L115M, C145A+S113C, and C145A+S10C. These double mutant variants had higher dimerization in the apo state than the corresponding single mutant network variants (but less than C145A M^pro^, which is 100% dimeric under the same conditions) (**Figure S4**). When the double mutant variants bound to the non-covalent compound X11612, the dimer fraction was similar to C145A M^pro^, with similar fractions of singly and doubly bound dimers. Additionally, the double mutant variants bound to substrate like C145A—about 50% apo, 30% singly bound, and less than 10% doubly bound dimers (**Figure S4**). Except the increased fraction of apo dimers for C145A+S113C, the double mutant variants displayed a dimerization behavior similar to C145A M^pro^; thus, the network mutations did not considerably impact substrate-induced dimerization of M^pro^.

### Heterodimerization of two inactive M^pro^ variants rescued enzymatic activity

Upon addition of excess quantities of C145A M^pro^ to ensure complete heterodimerization, the network variants displayed 2-to 79-fold activity rescue per monomer (**Figure 4b**). The most modest increase was for S10A, where the S10A/C145A heterodimer reached 2.4-fold activity of the starting homodimer. The variants with the greatest degree of activity boost compared to their respective homodimer were S10C (79-fold), S113C (42-fold), L115M (24-fold), and L115A (22-fold). Strikingly, while the homodimeric S113C variant had only 1% of WT activity, the S113/C145A heterodimer reached 50% of WT activity under the same assay conditions. Thus, quite unexpectedly, heterodimerization of two inactive monomers of M^pro^—one catalytically inactivated and one with defective cleavage—combined to result in rescue of enzymatic activity.

The catalytically inactivated network variants (double mutants) allowed us to tease out the contributions of the monomer in *cis*-versus *trans*-of substrate cleavage to the observed rescue of enzymatic activity. We generated two additional types of heterodimers with the double mutant variants: WT dimerized with a double mutant variant (WT/C145A+network mutation), and network variant dimerized with the catalytically inactivated double mutant counterpart (network mutation/C145A+network mutation).

In the heterodimers of WT with double mutant network variants (WT/C145A+network mutation), activity per monomer was rescued completely, to the level of WT/C145A heterodimer (**Figure 4c**). This result suggests that addition of the network mutation to the *trans* monomer (C145A+network mutation) does not affect *cis* cleavage by the WT monomer. In the network/double mutant heterodimers, cleavage rates were about 50-100% of WT homodimers. This indicates that the inactivated network mutant in *trans* of the cleaving active site can rescue the activity of the network variant monomer. Furthermore, in all cases we tested, this type of heterodimer (network mutation/C145A+network mutation) achieved more activity than the heterodimer of a network variant with C145A (network mutation/C145A); the network mutation in *trans* of the cleaving active site may even slightly improve the apparent activity, but at least *does not reduce* the heterodimer’s activity. In all cases, the network mutation in the *cis* (cleaving active site), but not the *trans*, monomer was debilitating to the enzymatic activity.

Thus, by systematically testing different heterodimeric combinations, we demonstrated that even a catalytically inactive network variant monomer can enhance cleavage activity in *cis*— whether the cleaving monomer is WT or contains a network mutation.

## Discussion

SARS-CoV-2 M^pro^ is active as an obligate dimer, but how one monomer affects the other and the functional consequences of cooperativity between the two monomers have remained unclear. In this work, we have probed the inter-monomer communication of M^pro^ through enzymology, mutagenesis, and structural biology. We found that the two active sites are connected through a structural network, and binding at one active site is communicated to the other. This means that the two monomers are functionally interdependent. Our results support a mechanism where the two active sites cooperate to mediate substrate cleavage by long-range (~40 Angstroms) allosteric communications.

The functional asymmetry and catalytic activation of one monomer by ligand binding at the *other* monomer have been revealed by our heterodimer assay results. Both substrate and inhibitor binding at the active site of *trans* monomer enhanced substrate cleavage in the *cis* monomer, indicating kinetic allostery. Introducing mutations to the residues in the putative network bridging the two active sites severely decreased activity, likely disrupting this allosteric communication. Mutations within the network, even distal from the active site and/or dimer interface, were disruptive to substrate cleavage. The impaired catalytic efficiency was largely driven by reduced turnover rate (*k*_cat_), and modest decrease in binding affinity. The network variants dimerized efficiently in the presence of ligand, suggesting that their catalytic defects stemmed from disrupted communication, not simply from poor dimerization. Interestingly, mutations to the network residues not in the *trans* but rather in the *cis* monomer were disruptive. Perhaps even more interestingly, heterodimerizing two catalytically defunct variants, C145A and a catalytically deficient network mutant, largely recovered enzymatic activity. This rescue could not be explained by improved dimerization alone and instead points to the functional asymmetry between the monomers. As previously suggested^13^, we present strong evidence for positive kinetic cooperativity, where productive turnover at one active site depends on substrate binding at the other. Our findings support a mechanism in which M^pro^ dimeric architecture enables long-range communication and cooperative substrate processing, explaining why dimerization is both necessary and functionally advantageous for this essential viral protease.

We propose a model for M^pro^’s reaction cycle consistent with our findings (**Figure 5**). M^pro^ monomer-dimer equilibrium shifts to dimers in the presence of substrate, in accordance with the presence of ligand-induced dimerization reported for M^pro^ and its homologs^12,32,34^. However, our results suggest that positive cooperativity in the observed enzymatic activity is not for binding (thermodynamic cooperativity) but rather the cleavage reaction (kinetic cooperativity). The considerable fraction of singly bound dimers suggests the lack of strong positive cooperativity in ligand binding. The lack of positive cooperativity in binding corroborates reports of negative cooperativity from other studies^13,17^. Once substrate binds to both active sites, inter-monomer communication primes the second active site for substrate cleavage. Our evidence here suggests *against* a model in which substrate cleavage by the first monomer induces cleavage at the second monomer –WT/C145A heterodimers would show reduced M^pro^ activity per monomer if this were the case.

**Figure 5.**
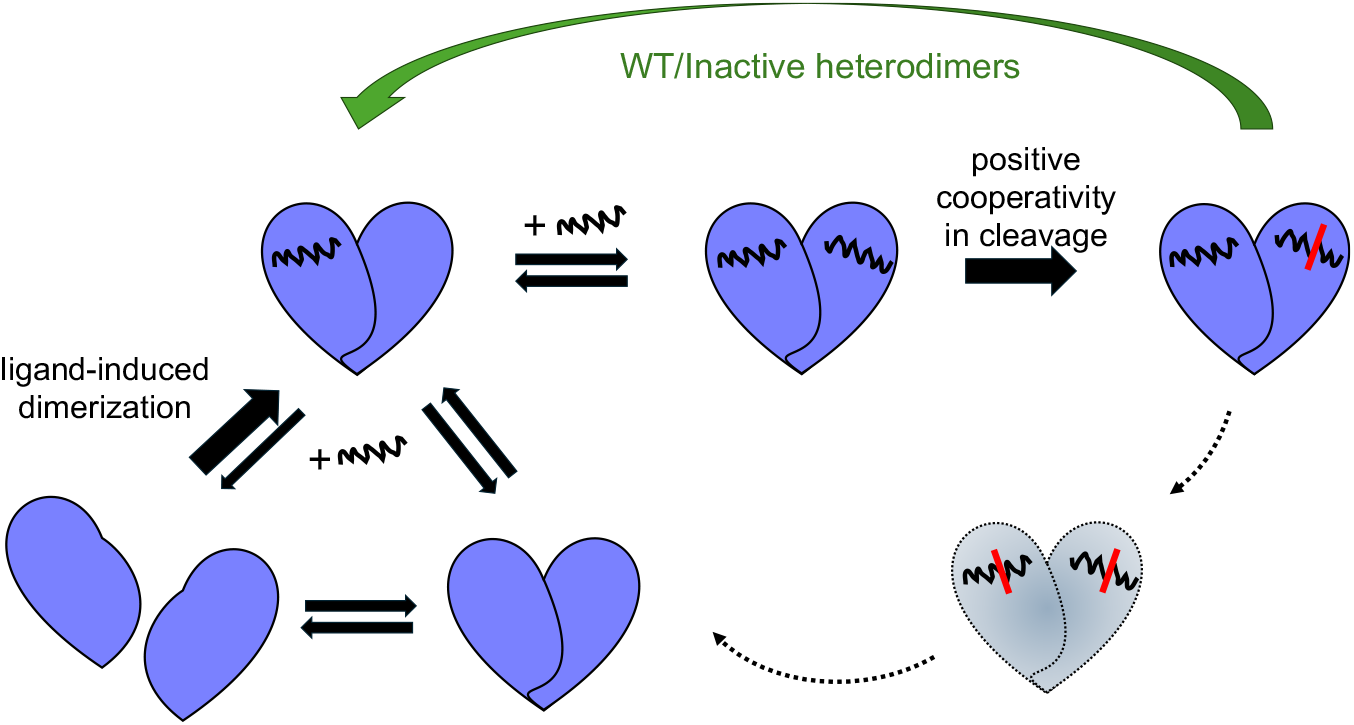
Proposed model for cooperative activity of SARS-CoV-2 M^pro^. The monomer-dimer equilibrium shifts to dimers in the presence of substrate (wiggly black line), and positive cooperativity induces cleavage of the second substrate upon binding (indicated by red line). Steps and intermediate(s) with no direct evidence are indicated by dashed lines. Product release and dissociation are likely to slow down the apparent rate of reaction relative to heterodimers, for which constitutive binding at the inactive monomer would speed up the apparent rate of reaction.

While the importance of dimerization in M^pro^’s catalytic activity is established, the asymmetry between the two monomers and any implications of this asymmetry for catalytic activity are still debated. We previously reported cocrystal structures of M^pro^ with a series of natural substrate peptides as well as inhibitor-bound structures for both SARS-CoV-2 and other divergent coronaviruses^19,23^. In all these structures, our careful analysis confirmed that both active sites were fully occupied with virtually identical protomers, which is *not* consistent with classical “half-site reactivity” mechanism^18^. Our heterodimer experiments with irreversible covalent inhibitors also refute this model. The asymmetry previously reported for apo SARS-CoV M^pro^ is likely due to crystal packing (one of the chain termini contacts the crystal symmetry mate), leading to the suggestion that apo dimers are asymmetric and only one protomer is in the “active conformation”^18^. However, in the apo state and in solution there is no direct evidence and no inherent reason why the two protomers would be asymmetric. In MD simulations with longer equilibration and sampling time, the apo and substrate-bound monomers behave similarly regardless of context, with the loops directly interacting with the substrate becoming less flexible when bound, as expected (**Table S2**). Our results and reaction cycle model we propose here are consistent with more recent reports suggesting that the two active sites are equivalent in the apo state and can be simultaneously functional;^9^ the asymmetry induced by the first substrate binding would be resolved after second substrate binding.

Multimerization and allostery are fundamental mechanisms through which biological molecules regulate function and activity. Dimerization and cooperativity in coronaviral M^pro^ are likely essential to fine tune the enzymatic activity in response to substrate concentration. The initial steps of the reaction cycle, namely substrate-induced dimerization and the requirement for both active sites to be occupied for cleavage, would ensure that viral polyprotein cleavage occurs only under specific intracellular conditions, particularly within the insulated replication organelle with sufficiently high M^pro^ and substrate concentrations. Our results highlight the essential role of allosteric communication between the two active sites in mediating cooperative activity and substrate cleavage to allow proper processing of the viral polyproteins.

## Materials and Methods

### Protein overexpression and purification

WT and variant M^pro^ proteins were expressed with an N-terminal polyhistidine-SUMO tag and purified as described previously^19,23,35,36^. The point mutations were introduced into the PETite expression plasmid using site-directed mutagenesis. Mutations were confirmed with plasmid sequencing and variant protein identities confirmed by characteristic molecular weight in native mass spectrometry.

### Protein crystallization

For cocrystallization, complexes with PF-00835231 were assembled by incubating 6 mg/mL of each M^pro^ with a 10-fold molar excess of inhibitor for 1 h at room temperature. The solutions were spun down at 10,000xg for 10 minutes to remove any insoluble compound or protein aggregates. Protein crystals were obtained with 18–22% (*w*/*v*) PEG 3350, 0.15-0.22 M NaCl, and 0.1 M Bis-Tris Methane pH 5.5 by hanging drop vapor diffusion at room-temperature in pre-greased VDX trays (Hampton Research, Aliso Viejo, CA, USA). Varying the protein-to-mother liquor ratios (1 µL:2 µL, 2 µL:2 µL, 3 µL:2 µL) helped obtain large, diffraction-quality crystals. To limit vibrational stress, crystallization trays were placed on foam padding. Cocrystals of M^pro^ variants with PF-00835231 appeared overnight and grew fully within 2 weeks.

### Crystallographic data collection and structure

Crystals were sent for data collection at the Brookhaven National Laboratory NSLS-II Beamline 17-ID-2 (FMX). X-ray diffraction data were collected at 100 K. Cocrystals were soaked in cryogenic solution made by supplementing the exact precipitant solution with 25% glycerol, then looped and frozen in liquid nitrogen. At NSLS-II, the collected diffraction intensities were automatically indexed, integrated, and scaled using XDS^37^. All structures were determined using molecular replacement with PHASER^38^. The reference model used was PDB 8DSU. Prior to molecular replacement, the model was modified by removing all water, buffer, and cryogenic molecules as well as the small molecule inhibitor in the active site. To minimize reference model bias, 5% of the data was reserved to calculate R_free_^39^. Before fitting inhibitor atoms into the electron density, inhibitor geometry was optimized in Gaussview 6 using Gaussian 16 with the basis set: DFT B3LYP 6-311++G (d,p). Model building and refinement were performed using Coot^40^ and Phenix^41^.

X-ray data collection parameters and refinement statistics are presented in Supplemental Material, Table S1.

### Structural analysis

Structural analyses and figure generation were carried out in PyMOL by Schrödinger, LLC^42^. Hydrogen bonds were identified using the show_contacts plugin with default setting, which define a hydrogen bond as having a bond angle between 63° and 180°, and a distance of less than 4.0 Å (or under 3.6 Å for an ideal hydrogen bond) between the proton donor and the acceptor heavy atom.

The SARS-CoV-2 M^pro^ substrate cocrystal structures published previously^19^ (7T70, 7T8M, 7T8R, 7T9Y, 7TA4, 7TA7, 7TB2, 7TBT, 7TC4) were analyzed by first removing the solvent and creating symmetry-mate if needed to obtain the dimeric structure. For each structure, the distance between all alpha-carbon atom pairs were calculated. These distances were then compared between two structures to calculate the “distance differences”. The average “distance difference” for each residue pair was calculated across all structures and ranked to find the most positionally consistent residues across all nine substrate cocrystal structures.

### Differential scanning fluorimetry (DSF)

Thermal shift assays were performed in final conditions of 2 µM M^pro^, 50 mM Tris pH 8.0, 300 mM NaCl, 2% DMSO, with 5x Sypro orange dye (Invitrogen 5000x Sypro), as optimized and described previously^25^. Thermal denaturation spectra were obtained in a Thermo Scientific AB-0700 96-well OCR plate using a Bio-Rad CFX Real Time PCR thermocycler. Temperatures during the melt ranged from 25 – 95 °C, with intervals of 0.3 °C every 12 seconds. The HEX channel was utilized to measure relative fluorescence units (RFU) for each well at each temperature. Data were analyzed as reported previously^25^.

### Enzyme activity assays

To determine the enzyme kinetic parameters of M^pro^ variants, 75 to 1000 nM enzyme was added to a diluted series of 0–200 μM FRET substrate (Dabcyl-KTSAVLQSGFRKM-Glu(Edans) (GenScript)) in assay buffer (50 mM Tris pH 7.5, 50 mM NaCl, 1 mM ethylenediaminetetraacetic acid (EDTA), 1 mM dithiothreitol (DTT), and 4% dimethyl sulfoxide (DMSO)) as described previously^25^. Other studies with sufficient data report WT SARS-CoV-2 M^pro^ Hill coefficients between 1.4 and 1.6^13–16^ which depended on assay conditions. We reproduced these values with low DMSO concentrations (data not shown), where the fluorogenic peptide displayed visible aggregation; therefore, all comparisons were conducted at 4% DMSO to ensure substrate solubility.

The cleavage reaction was monitored using a PerkinElmer EnVision plate reader at room temperature (340 nm excitation and 492 nm emission). Three or more replicates were performed for each variant. The initial velocities (RFU/s) were plotted against substrate concentrations and fit using GraphPad Prism 10^43^ to the Michaelis–Menten equation to calculate enzymatic parameters; the same data were also fit to the Hill equation to obtain allosteric sigmoidal fit parameters V_max_ and K_0.5_ (**Supp. Figure 2b**). RFU was converted to concentration using a linear fit of the calibration curve obtained with different concentrations of substrate and the end point of cleavage product with WT enzyme, which gave a factor of 6000 RFU per μM under the assay conditions.

### Native mass spectrometry

Each protein was purified in SEC buffer (25 mM HEPES pH 7.5, 150 mM NaCl, and 1 mM TCEP) and then diluted to 2-4 mg/mL in 50-100 µL in the same buffer. The protein was dialyzed for three hours into 200 mM ammonium acetate (pH 6.8) using a Slide-A-Lyzer MINI Dialysis Device (0.5 mL; MWCO 20 kDa; Thermo Fisher Scientific), then dialyzed again overnight. After dialysis, the protein concentration was re-determined based on absorbance at 280 nm. Each protein was diluted into ammonium acetate to 5 µM. Native mass-spectrometry analysis was performed on a QE UHMR mass spectrometer (Thermo Fisher Scientific) equipped with TriVersa NanoMate (Advion Interchim Scientific) with parameters as follows: Positive mode; spray voltage: 1.5 kV; resolution: 12,500; mass range (*m*/z): 2000-8000. The mass spectra were analyzed using PMI-Byos Intact software (Protein Metrics Inc.). Each protein was analyzed in triplicate.

For protein-ligand assessments, 2.5 mM ligand stock solution was prepared in DMSO; 5 µM protein solution was prepared in 200 mM ammonium acetate. 0.2 µL of the 2.5 mM ligand stock solution was transferred into 19.8 µL of the 5 µM protein solution. Protein and ligand were incubated at 1:5 molar ratio (5 µM : 25 µM) for 1 hour at room temperature, and then protein was dialyzed and analyzed as described above.

### Purification of irreversibly inhibited WT protein

An aliquot of concentrated WT SARS-CoV-2 M^pro^ stock (300 µM) was thawed at room temperature. The Michael acceptor, covalent peptidomimetic inhibitor N3 (Tocris, catalog no. 7230) was solubilized to 50 mM in 100% dimethyl sulfoxide (DMSO), then added to the M^pro^ solution at a final molar ratio of 10:1 (inhibitor:active site). The resulting protein–inhibitor mixture was incubated at room temperature overnight. Enzymatic activity assays were intermittently performed on small aliquots throughout the incubation to monitor the extent of M^pro^ inhibition. Complete inhibition was confirmed the following morning by the absence of detectable enzymatic activity even at [E] > 300 µM.

To remove unbound inhibitor and exchange the buffer, the protein–inhibitor solution was subjected to size-exclusion desalting using Zeba 0.5 mL spin columns with a 1,000 Da molecular weight cutoff (Thermo Scientific, catalog no. 89883). The sample was then dialyzed against assay buffer (50 mM Tris-HCl, pH 7.5; 50 mM NaCl; 1 mM ethylenediaminetetraacetic acid [EDTA]; 1 mM dithiothreitol [DTT]) for 4 hours, followed by a second dialysis step overnight in fresh buffer. The absence of free N3 in the final sample was verified by liquid chromatography–mass spectrometry (LC-MS). Final protein aliquots were flash-frozen in liquid nitrogen and stored at –80 °C.

### Heterodimer activity assays

To generate WT/C145A heterodimers, WT M^pro^ and catalytically inactive C145A M^pro^ were thawed on ice and then brought to room temperature (RT) once no visible ice remained. C145A M^pro^ was added to a final concentration of 5 µM into assay buffer (50 mM Tris-HCl, pH 7.5; 50 mM NaCl; 1 mM ethylenediaminetetraacetic acid [EDTA]; 1 mM dithiothreitol [DTT] followed by the addition of WT M^pro^ to a final concentration of 20 or 40 nM.

Network variant heterodimers were generated following the same protocol. In all cases, the inactive (including C145A mutation) variant was added first at a final concentration of 5 µM, followed by the active component to a final concentration of 20 nM. In one set of samples, 20 nM network variant was paired with 5 µM catalytically inactive C145A M^pro^. In a second set, 20 nM network variant was combined with 5 µM of a double mutant containing both the network variant and the C145A substitution. A third set of samples consisted of 20 nM WT M^pro^ as the active component and 5 µM of the double mutant as the inactive partner.

For samples containing only active protease (WT M^pro^ or network variant) bovine serum albumin (BSA; Sigma, catalog no. P5369) was added at a concentration equivalent in mass to that of 5 µM WT M^pro^, to control for molecular crowding and stabilization of dimerization. Samples were mixed immediately by inversion, and the time of mixing was designated as time zero.

Proteolytic activity was monitored using a fluorogenic FRET peptide substrate representing the nsp4–nsp5 cleavage junction, as previously described^25^. The final substrate concentration was 40 µM. Initial cleavage velocities were determined at multiple time points between 10 and 300 minutes; equilibration was reached by all samples at about 120 minutes. All measurements were performed in triplicate for each time point.

### Cleavage activity of semi-inhibited heterodimers

To evaluate the proteolytic activity of semi-inhibited M^pro^ heterodimers, several samples were assembled and tested in parallel. N3-inhibited WT M^pro^, purified as described in the section above, was used as the inactive species at a final concentration of 5 µM in relevant samples.

The active component, catalytically competent WT M^pro^, was included at either 20 nM or 40 nM, as specified below. The following combinations were prepared:

1. 20 nM WT M^pro^ + 5 µM C145A M^pro^
2. 40 nM WT M^pro^ + 5 µM C145A M^pro^
3. 5 µM N3-inhibited WT M^pro^ alone
4. 20 nM WT M^pro^ + 5 µM N3-inhibited WT M^pro^

Reactions were conducted in assay buffer (50 mM Tris-HCl, pH 7.5; 50 mM NaCl; 1 mM EDTA; 1 mM DTT). For each sample, the inactive component was added first, and the reaction was initiated by the addition of the active WT M^pro^, followed by immediate gentle inversion to mix. The moment of mixing was defined as time zero.

Initial cleavage velocities were measured using a fluorogenic FRET peptide substrate corresponding to the nsp4–nsp5 cleavage site (final concentration: 40 µM). Aliquots were tested at multiple time points over a 96-hour period. All measurements were performed in triplicate, and equilibrium was reached by approximately 72 hours.

## Acknowledgments

This research was supported by the National Institute of General Medical Sciences (NIGMS) R35GM151996 and from Novartis Institutes for Biomedical Research (NIBR). S.N.Z. was supported by National Institute of Allergy and Infectious Diseases (NIAID) F31AI179014.

