## Supplemental Material for "Cooperativity and Communication between the Active Sites of the Dimeric SARS-CoV-2 Main Protease"

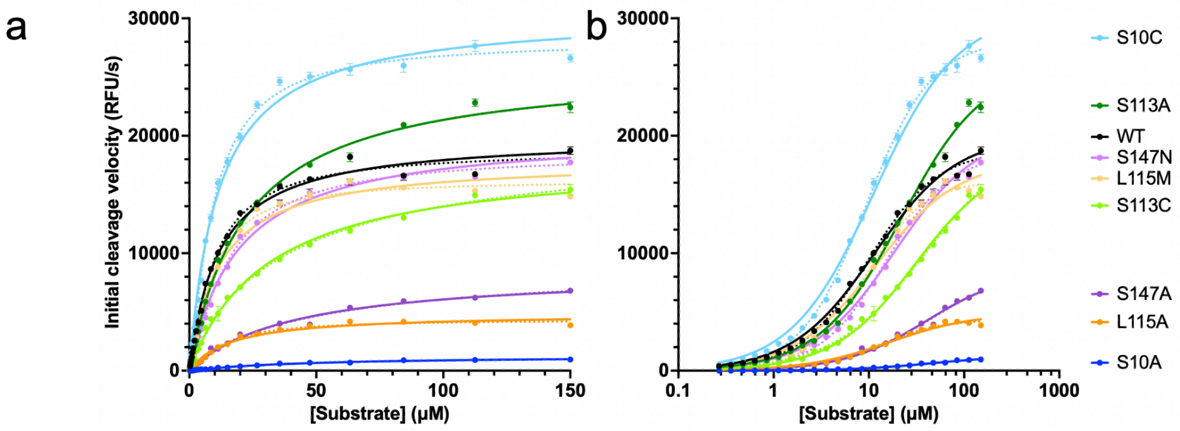


c

| $M^{pro}$ Variant | $K_m$ (μM) | $V_{max}$ (μM/s) | [E] (μM) | $k_{cat}$ (1/s) | $k_{cat}$ / $K_m$ 1/(μMs) | Allosteric fit $K_{0.5}$ (μM/s) | Allosteric fit $V_{max}$ (μM/s) | Hill Coefficient | Relative $k_{cat}$ / $K_m$ |
|---|---|---|---|---|---|---|---|---|---|
| WT | 11.4 ± 0.6 | 3.3 ± 0.05 | 0.1 | 33.2 ± 0.5 | 2.9 ± 0.2 | 9.8 ± 0.6 | 3.1 ± 0.7 | 1.22 ± 0.05 | 100% |
| S10A | 49.1 ± 4.6 | 0.2 ± 0.01 | 1.0 | 0.2 ± 0.01 | 0.004 ± 0.0005 | 38.8 ± 5.0 | 0.2 ± 0.01 | 1.13 ± 0.07 | 0% |
| S10C | 11.4 ± 0.8 | 5.1 ± 0.1 | 0.5 | 10.2 ± 0.2 | 0.9 ± 0.07 | 9.4 ± 0.3 | 4.7 ± 0.06 | 1.26 ± 0.04 | 30% |
| S113A | 22.2 ± 1.3 | 4.3 ± 0.09 | 0.5 | 8.7 ± 0.2 | 0.4 ± 0.02 | 22.0 ± 1.7 | 4.3 ± 0.1 | 1.00 ± 0.04 | 13% |
| S113C | 31.2 ± 2.3 | 3.1 ± 0.09 | 1.0 | 3.1 ± 0.1 | 0.1 ± 0.01 | 39.4 ± 3.8 | 3.3 ± 0.1 | 0.90 ± 0.03 | 3% |
| L115A | 16.4 ± 1.3 | 0.8 ± 0.02 | 0.63 | 1.3 ± 0.03 | 0.08 ± 0.01 | 12.6 ± 0.7 | 0.7 ± 0.02 | 1.36 ± 0.08 | 3% |
| L115M | 11.8 ± 0.7 | 3.0 ± 0.05 | 0.25 | 11.9 ± 0.2 | 1.0 ± 0.06 | 9.3 ± 0.4 | 2.7 ± 0.05 | 1.41 ± 0.07 | 35% |
| S147A | 40.7 ± 3.4 | 1.4 ± 0.05 | 1.0 | 1.4 ± 0.1 | 0.04 ± 0.003 | 47.4 ± 9.4 | 1.5 ± 0.1 | 0.94 ± 0.07 | 1% |
| S147N | 18.9 ± 1.0 | 3.4 ± 0.06 | 1.0 | 3.4 ± 0.1 | 0.2 ± 0.01 | 15.0 ± 0.7 | 3.1 ± 0.05 | 1.24 ± 0.05 | 6% |

**Figure S1**: Enzymatic parameters for WT SARS-CoV-2 $M^{pro}$ and network variants. (**a**) The initial rate of cleavage versus substrate concentration for each network variant, fit to the Michaelis-Menten and allosteric sigmoidal equations to yield parameters. Solid and dashed lines show Michaelis-Menten and allosteric sigmoidal fit respectively. (The enzyme concentration was adjusted for each variant to enable sufficient detectable activity; see column [E] in panel c). **b**) The same data on a log10 scale. (**c**) Enzyme kinetic parameters obtained from the fits in panels (a and b): substrate affinity ($K_m$), maximum velocity ($V_{max}$), concentration of enzyme used for the assay ([E]), catalytic efficiency ($k_{cat}/K_m$), and catalytic efficiency relative to the WT enzyme. $K_{0.5}$, $V_{max}$, and Hill coefficients were also calculated from an allosteric sigmoidal fit.

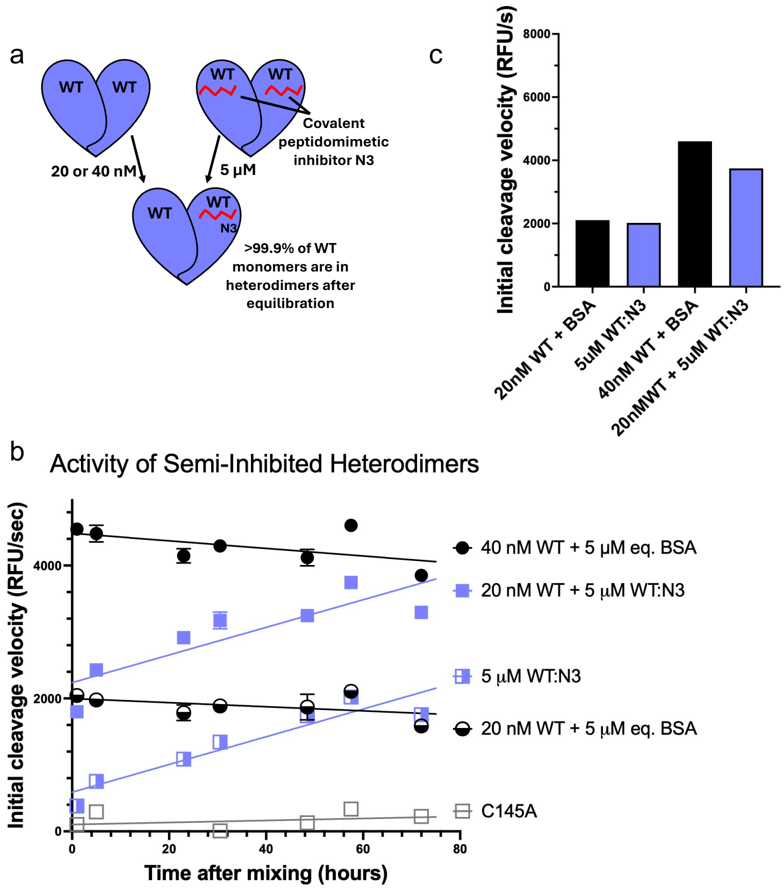


**Figure S2**: Mixing WT M$^{pro}$ with inhibited inactive WT:N3 resulted in increase in apparent enzymatic activity. **a**) A relatively small amount of WT M$^{pro}$ (20 or 40 nM) was added to an excess amount of purified M$^{pro}$ bound to the covalent irreversible inhibitor N3 (WT:N3). **b**) The enzymatic activity assessed by initial cleavage velocity of substrate peptide over time for WT homodimers and for semi-inhibited heterodimers (WT/WT:N3). WT:N3 species increased in activity which can be attributed to trace amount (<0.5%) of WT heterodimerizing with inhibited monomers. **c**) The enzymatic activity of WT M$^{pro}$ and heterodimers with inhibited inactive WT:N3 after equilibration.

a

| $M^{pro}$ variant | Apo $T_M$ ± SEM (°C) | Percent dimer in apo | Percent dimer bound to S | Percent dimer bound to non-covalent inhibitor |
|---|---|---|---|---|
| WT | 52.9 ± 0.3 | 69 ± 3 | N.D. | 99 ± 0 |
| C145A | 56.8 ± 0.1 | 98 ± 1 | 99 ± 1 | 100 ± 0.3 |
| S10A | 51.7 ± 0.1 | 9 ± 2 | N.D. | 47 ± 3 |
| S10C | 52.1 ± 0.1 | 12 ± 1 | 92 ± 1* | 76 ± 0.3 |
| S113A | 49.8 ± 0.05 | 14 ± 1 | N.D. | N.D. |
| S113C | 51.2 ± 0.1 | 10 ± 3 | 63 ± 3* | 83 ± 2 |
| L115A | 47.5 ± 0.03 | 9 ± 1 | N.D. | 47 ± 4 |
| L115M | 50.2 ± 0.1 | 15 ± 1 | 97 ± 0.3* | 87 ± 1 |
| S147A | 51.2 ± 0.2 | 7 ± 1 | N.D. | 48 ± 5 |
| S147N | 47.8 ± 0.05 | 8 ± 1 | N.D. | 67 ± 1 |

* These samples are C145A + network mutation, rather than just the network mutation.

b

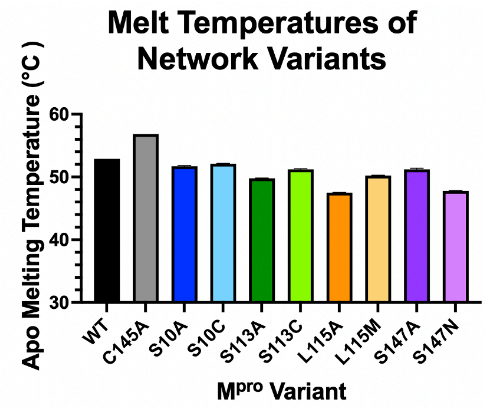


**Figure S3**: (a) Apo melting temperatures, apo dimerization, substrate-bound dimerization, and inhibitor-bound dimerization for each Mpro variant. Thermostability assays were performed with 2 μM of protein. Dimerization experiments were performed with 5 μM of protein and 25 μM of ligand. Each value results from three replicate experiments. (b) Bar graph comparing apo thermostability for each variant.

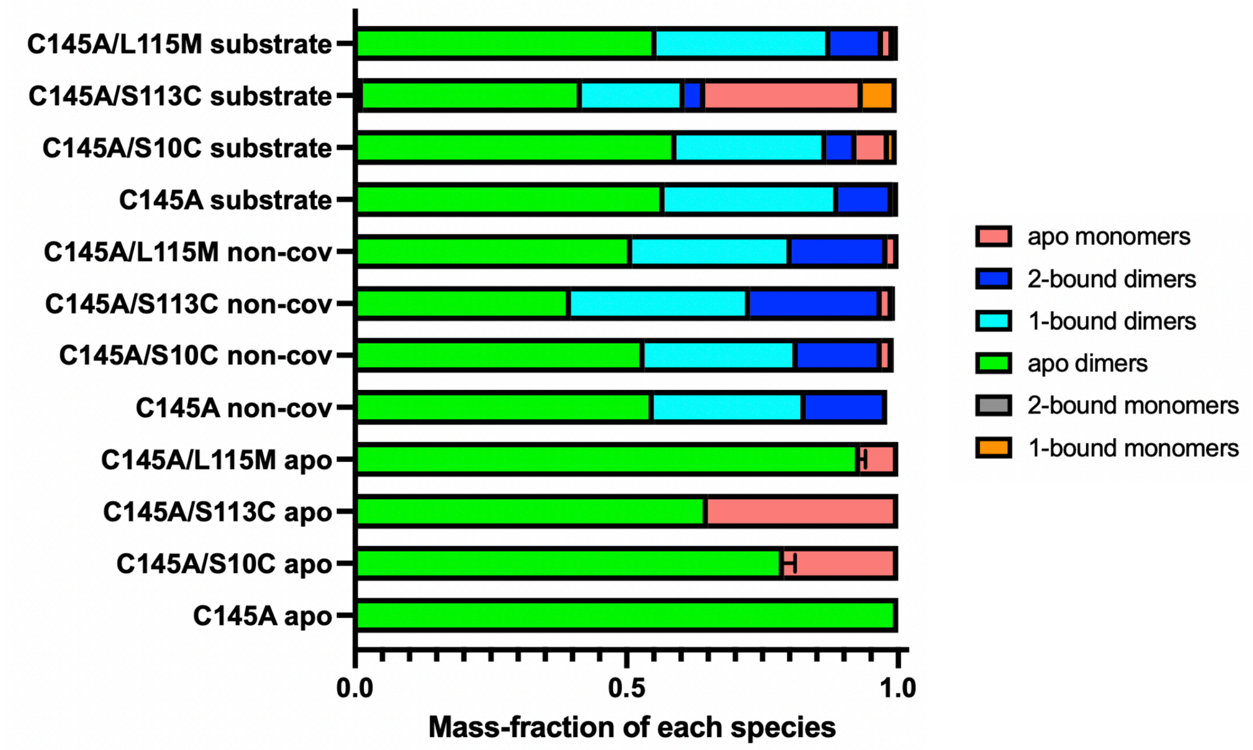


**Figure S4.** The fraction of monomeric and dimeric states for different inactive Mpro variants, analyzed by native mass spectrometry. The non-covalent inhibitor (Moonshot X11612) and substrate (peptide sequence representing nsp4-nsp5) were used as ligands. Error bars indicate standard error of the mean from three replicates.

**Table S1.** Crystallization and structure refinement statistics of SARS-CoV-2 $M^{pro}$ variants S10A, S10C, S113A, S113C, L115A, L115M, S147A, and S147N in complex with PF-00835231.

| **$M^{pro}$ Variant** | **S10A** | **S10C** | **S113A** | **S113C** | **L115A** | **L115M** | **S147A** | **S147N** |
|---|---|---|---|---|---|---|---|---|
| **PDB ID** | 9O6D | 9O6E | 9O6F | 9O6P | 9O6Q | 9O74 | 9OPM | 9OPN |
| **DATA COLLECTION** | | | | | | | | |
| Location | NSLS II | NSLS II | NSLS II | NSLS II | NSLS II | NSLS II | NSLS II | NSLS II |
| Resolution Range (Å) | 33.61-1.90 (1.97-1.90) | 33.72-1.99 (2.06-1.99) | 31.93-1.97 (2.04-1.97) | 33.66-2.28 (2.36-2.28) | 33.56-2.21 (2.29-2.21) | 33.68-2.19 (2.27-2.19) | 33.55-2.13 (2.21-2.13) | 33.57-1.77 (1.83-1.77) |
| Space Group | P 1 21 1 | P 1 21 1 | P 1 21 1 | P 1 21 1 | P 1 21 1 | P 1 21 1 | P 1 21 1 | P1 21 1 |
| a,b,c, (Å) | 54.84, 98.91, 59.03 | 54.97, 99.39, 59.36 | 55.10, 99.33, 59.62 | 54.39, 98.57, 58.61 | 54.71, 98.97, 58.83 | 54.78, 99.36, 59.11 | 55.02, 98.63, 58.64 | 55.05, 98.48, 58.49 |
| alpha, beta, gamma (°) | 90,107.56, 90 | 90, 107.55, 90 | 90, 107.84, 90 | 90, 106.72, 90 | 90, 107.52, 90 | 90, 107.54, 90 | 90, 107.64, 90 | 90, 107.39, 90 |
| Total Reflections | 94,180 | 83,307 | 84,593 | 52,868 | 58,073 | 61,090 | 66, 214 | 111,846 |
| Unique Reflections | 47,166 | 41,682 | 42,646 | 26,854 | 29,455 | 30,548 | 33,226 | 56,403 |
| Multiplicity | 2.0 (2.0) | 2.0 (2.0) | 2.0 (2.0) | 2.0 (2.0) | 2.0 (2.0) | 2.0 (2.0) | 2.0 (2.0) | 2.0 (2.0) |
| Completeness (%) | 99.81 (99.51) | 100 (100) | 98.8 (98.7) | 99.0 (99.8) | 98.3 (98.2) | 98.4 (97.6) | 99.5 (99.1) | 97.5 (96.5) |
| Average I/sigma(I) | 16.3 (3.1) | 12.5 (2.6) | 16.5 (3.9) | 14.6 (3.9) | 9.5 (3.0) | 10.9 (3.4) | 8.9 (3.1) | 14.2 (3.2) |
| Wilson B-Factor | 29.64 | 33.04 | 23.52 | 40.25 | 33.91 | 36.39 | 30.56 | 25.87 |
| $R_{merge}$ | 0.024 (0.223) | 0.03 (0.283) | 0.032 (0.187) | 0.027 (0.185) | 0.045 (0.246) | 0.04 (0.201) | 0.049 (0.230) | 0.028 (0.225) |
| CC1/2 | 0.999 (0.916) | 0.999 (0.811) | 0.998 (0.918) | 0.999 (0.944) | 0.997 (0.844) | 0.997 (0.913) | 0.996 (0.894) | 0.999 (0.909) |
| **REFINEMENT** | | | | | | | | |
| $R_{work}$ | 0.19(0.27) | 0.18 (0.24) | 0.2 (0.27) | 0.21 (0.28) | 0.18 (0.23) | 0.18 (0.24) | 0.17 (0.22) | 0.18 (0.24) |
| $R_{free}$ | 0.24 (0.34) | 0.22 (0.30) | 0.25 (0.36) | 0.27 (0.40) | 0.24 (0.31) | 0.24 (0.31) | 0.22 (0.28) | 0.21 (0.32) |
| **RMSD in:** | | | | | | | | |
| Bond Lengths (Å) | 0.01 | 0.02 | 0.006 | 0.002 | 0.002 | 0.006 | 0.005 | 0.007 |
| Bond Angles (°) | 1.01 | 1.3 | 0.75 | 0.48 | 0.57 | 0.67 | 0.93 | 0.97 |
| **Ramachandran:** | | | | | | | | |
| Favored (%) | 97.35 | 97.35 | 96.84 | 95.86 | 96.85 | 97.02 | 98.18 | 98.51 |
| Allowed (%) | 2.65 | 2.48 | 3.16 | 4.14 | 3.15 | 2.98 | 1.82 | 1.49 |
| Outliers (%) | 0 | 0.17 | 0 | 0 | 0 | 0 | 0 | 0 |
| Rotamer outliers (%) | 2.12 | 1.35 | 1.75 | 1.38 | 0.98 | 1.98 | 1.36 | 0.78 |
| **B-Factors:** | | | | | | | | |
| Average | 36.28 | 37.75 | 29.68 | 48.87 | 38.05 | 41.19 | 34.76 | 32.36 |
| Macromolecules | 35.43 | 37.20 | 28.97 | 48.77 | 37.45 | 40.81 | 34.01 | 31.44 |
| Solvent | 45.18 | 45.16 | 36.69 | 51.78 | 44.86 | 47.18 | 42.82 | 42.25 |

**Table S2.** Stability and flexibility of $M^{pro}$ monomers in apo, one-substrate bound and two-substrate bound states from molecular dynamics simulations, assessed by root-mean-squared fluctuation (RMSF) and root-mean-squared deviation (RMSD) of backbone Carbon-alpha atoms. The chain B (*) was bound with a substrate in the one-bound state.

| | Apo | One-Bound | Two-Bound |
|---|---|---|---|
| **RMSF** (Å) | | | |
| **Chain A** | 1.15 ± 0.08 | 0.97 ± 0.07 | 0.85 ± 0.01 |
| **Chain B*** | 1.03 ± 0.10 | 0.84 ± 0.07 | 0.84 ± 0.01 |
| **RMSD** (Å) | | | |
| **Chain A** | 1.75 ± 0.25 | 1.52 ± 0.27 | 1.23 ± 0.25 |
| **Chain B*** | 1.73 ± 0.34 | 1.14 ± 0.16 | 1.15 ± 0.34 |

### Supplemental Methods

#### Molecular Dynamics (MD) Simulations

The $M^{Pro}$ dimer structure bound to two substrates (PDB ID: 7T70[1]) was used as the starting point. Missing C-terminal residues (301–306 on chains A and B) were modeled using COOT and Phenix[2] with updated electron densities. Singly bound and apo complexes were generated by removing one or both substrates in PyMOL. Protonation states were assigned, and missing atoms added using PlayMolecule ProteinPrepare[3].

MD simulations were performed with AMBER 22[4] using the pmemd.CUDA[5] engine and the ff19SB force field[6] with OPC water and ion parameters[7]. Systems were solvated in a truncated octahedral box with a 10 Å buffer and neutralized with 68 $Na^+$ and 68 $Cl^-$ ions to achieve 0.15 M salt concentration. System setup was completed with tleap. Minimization and equilibration were conducted in nine steps, including restrained minimizations, heating from 100 K to 298 K, and gradual restraint reduction on solute atoms. Production MD was run for 2000 ns at 1 bar and 298 K, with the first 500 ns discarded as equilibration. SHAKE was applied to constrain bonds involving hydrogen, and a 12 Å cutoff was used for non-bonded interactions. Each system was simulated in triplicate for a total of 4.5 µs. Trajectory analysis was performed using cpptraj[8] to compute RMSD and RMSF of protein Cα atoms except for the six C terminal residues which were highly variable, and inhibitor heavy atoms at 20 ps intervals.

1. Shaqra AM, Zvornicanin SN, Huang QYJ, Lockbaum GJ, Knapp M, Tandeske L, Bakan DT, Flynn J, Bolon DNA, Moquin S, Dovala D, Kurt Yilmaz N, Schiffer CA. Defining the substrate envelope of SARS-CoV-2 main protease to predict and avoid drug resistance. Nat Commun. 2022;13(1):3556. Epub 20220621. doi: 10.1038/s41467-022-31210-w. PubMed PMID: 35729165; PMCID: PMC9211792.
2. Liebschner D, Afonine PV, Baker ML, Bunkoczi G, Chen VB, Croll TI, Hintze B, Hung L-W, Jain S, McCoy AJ, Moriarty NW, Oeffner RD, Poon BK, Prisant MG, Read RJ, Richardson JS, Richardson DC, Sammito MD, Sobolev OV, Stockwell DH, Terwilliger TC, Urzhumtsev AG, Videau LL, Williams CJ, Adams PD. Macromolecular structure determination using X-rays, neutrons and electrons: recent developments in Phenix. Acta Crystallographica Section D. 2019;75(10):861-77. doi: doi:10.1107/S2059798319011471.
3. Varela-Rial A, Maryanow I, Majewski M, Doerr S, Schapin N, Jimenez-Luna J, De Fabritiis G. PlayMolecule Glimpse: Understanding Protein-Ligand Property Predictions with Interpretable Neural Networks. J Chem Inf Model. 2022;62(2):225-31. Epub 20220103. doi: 10.1021/acs.jcim.1c00691. PubMed PMID: 34978201; PMCID: PMC8790755.
4. Case D, Aktulga HM, Belfon K, Ben-Shalom I, Brozell S, Cerutti D, Cheatham T, Cisneros GA, Cruzeiro V, Darden T, Duke R, Giambasu G, Gilson M, Gohlke H, Götz A, Harris R, Izadi S, Izmailov S, Kollman P. Amber 20222022.
5. Gotz AW, Williamson MJ, Xu D, Poole D, Le Grand S, Walker RC. Routine Microsecond Molecular Dynamics Simulations with AMBER on GPUs. 1. Generalized Born. J Chem Theory Comput. 2012;8(5):1542-55. Epub 20120326. doi: 10.1021/ct200909j. PubMed PMID: 22582031; PMCID: PMC3348677.
6. Tian C, Kasavajhala K, Belfon KAA, Raguette L, Huang H, Migues AN, Bickel J, Wang Y, Pincay J, Wu Q, Simmerling C. ff19SB: Amino-Acid-Specific Protein Backbone Parameters Trained against Quantum Mechanics Energy Surfaces in Solution. Journal of Chemical Theory and Computation. 2020;16(1):528-52. doi: 10.1021/acs.jctc.9b00591.
7. Li Z, Song LF, Li P, Merz KM, Jr. Systematic Parametrization of Divalent Metal Ions for the OPC3, OPC, TIP3P-FB, and TIP4P-FB Water Models. Journal of Chemical Theory and Computation. 2020;16(7):4429-42. doi: 10.1021/acs.jctc.0c00194.
8. Roe DR, Cheatham TE, 3rd. PTRAJ and CPPTRAJ: Software for Processing and Analysis of Molecular Dynamics Trajectory Data. J Chem Theory Comput. 2013;9(7):3084-95. Epub 20130625. doi: 10.1021/ct400341p. PubMed PMID: 26583988.
